# The genome sequence of the Jean-Talon strain, an archeological tetraploid beer yeast from Québec

**DOI:** 10.1101/2020.02.11.944405

**Authors:** Anna Fijarczyk, Mathieu Hénault, Souhir Marsit, Guillaume Charron, Tobias Fischborn, Luc Nicole-Labrie, Christian R Landry

## Abstract

The genome sequences of archeological yeast isolates can reveal insights about the history of human baking, brewing and winemaking activities and migration around the globe. A yeast strain called Jean-Talon was recently isolated from the vaults of the Intendant’s Palace of Nouvelle France on a historical site in Québec City. This site has been occupied by various breweries, starting from the end of the 17th century and until the middle of the 20th century. We sequenced the genome of the Jean-Talon strain with short and long reads and reanalyzed the genomes of hundreds of yeast strains to identify its species of origin and determine how it relates to other domesticated and wild strains. The Jean-Talon strain is a tetraploid strain with numerous aneuploidies, is partially sterile and most closely related to beer strains from the beer and bakery genetic groups and industrial strains from the United Kingdom and Belgium. We conclude from this that the Jean-Talon strain most likely derives from recent brewing activities that took place in the same location and not from wild yeast that could have been domesticated by the original brewers of the Nouvelle France on the site.

## Introduction

The budding yeast *Saccharomyces cerevisiae* has a long history of domestication by humans for the production of fermented food and beverages (Marsit *et al.* 2017). Among the oldest evidence of this domestication are traces of DNA from more than 3000 years old wine jars in Egypt (Cavalieri et al. 2003). Signs of beer and chemical traces of beer production dating back from the last half of the fourth millennium before Christ were identified in sumerian artifacts (Michel *et al.* 1992). Alcoholic beverages were also present in prehistoric China about 9000 years before present (McGovern *et al.* 2004). Human populations brought fermented produces with them during their migration, including wine (Marsit *et al.* 2017), coffee and cacao beans (Ludlow *et al.* 2016), and this human assisted migration has contributed to the genetic organization of today’s population structure of *S. cerevisiae* (Peter *et al.* 2018). For instance, the genome analysis of some ale beer strains recently revealed that they originated from both European wine strains and Asian rice wine strains (Fay *et al.* 2019).

The production of fermented beverages by European settlers in North America started early in the colonies, for instance in the North American territory that became Nouvelle France and eventually French Canada. A common drink for French settlers in the 17th century called “bouillon” was made of bread leaven that was incubated in water, producing a lightly alcoholic beverage. Artisanal and domestic beer brewing most likely started during that period (Moussette 1992). Later in the 17th Century, the first industrial brewery was founded by the French King representative, the Intendant Jean Talon (Moussette 1996).

To establish his brewery, Talon acquired a relatively large lot, at the crossing of St. Charles and St. Lawrence rivers in Québec City, and built a massive 40 meters long building with modern brewing equipment (Moussette 1994). The brewery was short lived (1670-1675). However, after the British conquest (1763), entrepreneurs turned to brewing again. The lot for Talon’s old brewery had been turned into a powder storage and the Intendant’s Palace (1686-1713), which was destroyed by a fire and served as warehouses for the remainder of Nouvelle France.

In the early 19th century, the site went back to the brewing industry, thanks to the most successful family in Québec City’s brewing business, the Boswells. Joseph Knight Boswell was an Irish migrant born in 1812 who had been trained in Scotland. He settled in Québec City in 1830 and quickly worked as brewmaster before opening his own business, Anchor Brewery, in the 1840’s. Boswell expanded his business throughout the 1840’s and finally linked his booming brewery with Talon’s brewery in 1852 by renting the plot on which Talon’s brewery was built in the 17^th^ Century (Guimont 1987). The French vaults that Boswell rented were not from Talon’s brewery, but from the Second’s Intendant’s Palace (1713-1725), built slightly north of the brewery’s original site. Boswell will first use the 18^th^ Century vaults to store beer. Then, in 1875, he had a large building built on top of the vaults for his malting equipment and operations (Fiset 2001). In the 1930’s the Brewery opened “Les Voûtes Talon”, a pub located in the historical vaults. Beer production came to an end after the events of 1966 where beer brewed in Québec City caused nearly 50 deaths in heavy drinkers over the course of a few weeks. Public health authorities linked this to the use of cobalt sulfate as a stabilizer in the brewing process (Morin and Daniel 1967). The public relations disaster truly killed the brand brewed in Québec City (then labeled “Dow”) and lead to the complete shut down, in 1968, of the facility located on the same lot as Jean Talon’s brewery, ending a nearly 300 years long history.

Here, we sequenced the genome of a yeast strain (Jean-Talon) that was isolated from the Second’s Intendants Palace’s vaults. We report the whole-genome sequence using short and long read sequencing and the comparative analysis of this genome with other sequenced genomes. Our results reveal that the strain is polyploid, partially sterile and harbors multiple aneuploidies. A phylogenetic analysis reveals that it is indeed a beer strain that is closely related to the other industrial beer and baking strains, suggesting that its origin dates from recent industrial activities on the site and not from earlier brewing activities.

## Materials and Methods

### Strain sampling

30 Yeast Mold Agar (YM) plates (Difco 271210) with 25ppm of chloramphenicol to prevent bacterial growth were prepared. 3 YM plates each were placed at 10 different locations in the vaults of the Intendant’s Palace of Nouvelle France in September 2010. At each location, the 3 YM plates were exposed to the environment for 10, 20 and 40 minutes and then incubated at 28 °C for 2 days. The yeast strain was banked at the Siebel culture collection as BRY # 480 and sent to the Landry laboratory for analysis in 2019.

### DNA content and ploidy

Measurement of DNA content was performed using flow cytometry and the SYTOX™ green staining assay (Thermo Fisher, Waltham, USA) as done in (Charron *et al.* 2019). Cells were first thawed from glycerol stock and streaked on solid YPD in 6 petri dishes (room temperature, 3 days) to have isolated colonies. The strain BY4742 (haploid) and MG009 (BY4741xBY4742) (diploid) were used as controls. Liquid YPD cultures of 1 ml from 90 Jean-Talon isolated colonies and the two controls in 96-deepwell (2ml) plates were inoculated and incubated for 24 h at room temperature. Multiple colonies were considered to account for the possibility of an unstable ploidy. Cells were subsequently prepared as in (Gerstein *et al.* 2006). Cells were first fixed in 70% ethanol for at least 1 h at room temperature. RNAs were eliminated from fixed cells using 0.25 mg/ml of RNAse A overnight at 37°C. Cells were subsequently washed twice using sodium citrate (50mM, pH7) and stained with a final SYTOX™ green concentration of 0.6 μM for a minimum of 1 h at room temperature in the dark. The volume of cells was adjusted to be around a cell concentration of less than 500 cells/μL. Five thousand cells for each sample were analyzed on a Guava^®^ easyCyte 8HT flow cytometer using a sample tray for 96-well microplates. Cells were excited with the blue laser at 488 nm and fluorescence was collected with a green fluorescence detection channel (peak at 512 nm). The distributions of the green fluorescence values were processed to find the two main density peaks, which correspond to the two cell populations, respectively, in G1 and G2 phases. The data was analyzed using R version 3.4.159.

### Sporulation and dissection

The frozen stock of the Jean-Talon strain was streaked for single colonies onto a fresh YPD agar plate (1% yeast extract, 2% glucose, 2% peptone, 2% agar). Three independent colonies were picked, and the cells were patched on a solid sporulation medium (1% Potassium acetate, 0.1% Yeast extract, 0.05% Glucose, 0.01% sporulation dropout, 2% Agar). The sporulation dropout was composed of 0.0125 g/L Histidine, 0.0625g/L Leucine, 0.0125g/L Lysine and 0.0125g/L Uracil. After 7 days of incubation at room temperature, for each patch, a lump of cells was picked with a 200 μL micropipette tip and resuspended into 100 μL of a zymolyase solution (4U/ml of Zymolyase, Zymolyase 20T, Bioshop Canada). After 20 minutes, cells were centrifuged for 20 seconds at 16,100 g and the zymolyase solution was removed and replaced with 100 μL of a 1M sorbitol solution. For each of the initial colonies, 24 tetrads were dissected on fresh YPD plates with a SporePlay™ dissection microscope (Singer Instruments, Somerset, UK). After 5 days of incubation at room temperature, plates were photographed, and fertility was determined as the number of visible colonies to the naked eye.

### Short-read library construction and sequencing

Genomic DNA was extracted from overnight culture derived from an isolated colony following standard protocols (QIAGEN DNAeasy, Hilden, Germany). The library was prepared with the Illumina Nextera kit (Illumina, San Diego, USA) following the manufacturer’s protocol and modifications described in (Baym *et al.* 2015). The library was sequenced with the 150 bp PE mode in a lane of HiSeqX (Illumina, San Diego, USA) at the Genome Quebec Innovation Center (Montréal, Canada). Genome-wide coverage reached 75x after duplicate reads removal. Raw sequencing reads are available at NCBI (PRJNA604588).

### Long-read library construction and sequencing

DNA was extracted following a standard phenol-chloroform method from an overnight culture inoculated with an isolated colony of the Jean-Talon strain. PCR-free libraries for Oxford Nanopore Technologies (ONT) sequencing were prepared (in multiplex with other yeast strains) with kits SQK-LSK109 and EXP-NBD104 (Oxford Nanopore, Oxford, UK). Sequencing was performed on a FLO-MIN106 (revC) flowcell on a MinION sequencer (MIN-101B) driven by a MinIT computer (MNT-001) running the MinKNOW software v3.3.2. Basecalling was performed on the MinIT with guppy v3.0.3. Demultiplexing was performed using the guppy_basecaller utility v3.1.5. Basecalled, demultiplexed reads are available at NCBI (PRJNA604588).

### Genotyping of the Jean-Talon strain

Illumina reads were mapped onto the S288C *S. cerevisiae* reference genome vR64.2.1 using bwa mem v0.7.17 (Li 2013). Duplicated reads were tagged using picard tools v2.18 (http://broadinstitute.github.io/picard/). Genotypes were called with GATK v3.8 (DePristo *et al.* 2011) using the HaplotypeCaller module with an option *-ERC BP_RESOLUTION* and GenotypeGVCFs module with an option *--includeNonVariantSites*, with assumed ploidy 2. Single nucleotide polymorphisms (SNPs) were filtered with VariantFiltration module, excluding variants annotated with QualbyDepth < 2, MappingQuality < 40, MappingQualityRankSumTest < −12.5, FisherStrand > 60, StrandOddsRatio > 3 and ReadPosRankSum < −8.0. Additionally, genotypes with quality < 20 (both GQ and RGQ) and coverage < 10 reads were masked. Indels were excluded.

### Combining Jean-Talon SNPs with other datasets

To combine variants of the Jean-Talon strain with the published yeast variants, VCF files with SNPs from (Fay *et al.* 2019) (hereafter “Fay et al. dataset”), and (Peter *et al.* 2018) (hereafter “1000 yeast dataset”) were downloaded. All positions in the Jean-Talon file were filtered to keep those present in the Fay et al. and 1000 yeast datasets using bcftools v1.9 (Li *et al.* 2009), after adjusting chromosome names. The two datasets were combined with the Jean-Talon VCF file separately using GATK v3.8 (DePristo *et al.* 2011) CombineVariants module with an option *-genotypeMergeOptions UNIQUIFY.* Multiallelic SNPs were removed from respective merged datasets using bcftools v1.9. Principal Component Analysis was performed using *SNPrelate* (Zheng *et al.* 2012) package in R v3.6.1.

### Genotyping and comparison of beer strains

To find yeast strains that are genetically closest to the Jean-Talon strain, we downloaded and mapped yeast genomes from genetic beer groups from 4 studies: (Gallone *et al.* 2016; Gonçalves *et al.* 2016; Peter *et al.* 2018; Fay *et al.* 2019) (Supplementary Table 1). In total we analyzed 319 strains, including the Jean-Talon strain. Reads from these strains were trimmed for the common Illumina adapters with Trimmomatic v0.36 (Bolger *et al.* 2014), and mapped to the S288C *S. cerevisiae* genome using bwa mem v0.7.17 (Li 2013). Duplicate reads were tagged with picard tools v2.18. SNPs were called and filtered with GATK v4.1, as described above, but excluding filters, which are affected by single end reads, such as FisherStrand and StrandOddsRatio. We retained only SNPs with less than 10% of missing data across all strains. The *SNPrelate* package (Zheng *et al.* 2012) in v3.6.1 was used to calculate identity by state and identity by descent. Neighbor-joining tree was built using identity by state matrix with package *ape*. Kinship coefficient matrix (identity by descent) was estimated with KING method of moment. To estimate nucleotide diversity and divergence between closely related strains, we genotyped all genomic positions in 5 strains closely related to the Jean-Talon strain (A.Muntons, A.S-33, BE005, CFI and CFN; A.Windson was not included due to high amount of missing data), three other beer strains from the Beer/baking group (CHK, CFP, A.T-58) and one strain from the Ale2 group (A.2565) using GATK v4.1, as described above but with ploidy *4n* (except for CHK which is diploid). Genotypes passing all the filters were transferred on four (or two) reference genome sequences using seqtk v1.3 (https://github.com/lh3/seqtk), separately for each strain, and other positions were marked as missing data. Sequences of 6041 single exon, non-overlapping genes were extracted to generate multiple sequence alignments with concatenated gene sequences using bedtools v2.25 (Quinlan and Hall 2010) and custom python v3.6.8 scripts. Diversity statistics and number of synonymous sites were calculated using mstatspop v.0.1beta (https://github.com/CRAGENOMICA/mstatspop). To estimate the number of generations separating Jean-Talon and its closely related strains, we calculated time of divergence with a related strain, relative to the divergence time with S288C reference (Green *et al.* 2006; Skoglund *et al.* 2011). To estimate proportion of the branch length after split of Jean-Talon with the relative, we counted synonymous variants shared between Jean-Talon and S288C, but not with the relative, and variants shared between relative and the outgroup, but not with the Jean-Talon, and took the average and divided by the total number of synonymous sites. Shared synonymous variants were identified using custom python v3.6.8 script after annotating variants in VCF file using SnpEff (Cingolani *et al.* 2012). Divergence time with S288C was estimated with molecular clock, assuming mutation rate 1.67E-10 (Zhu *et al.* 2014), and considering only synonymous substitutions.

We obtained copy number profiles in 250 bp non-overlapping windows and in all genes with Control-FREEC v11.5 (Boeva *et al.* 2011, 2012). First, we estimated ploidy of each strain using nQuire using reads with mapping quality > 30, and *lrdmodel* option (Weiß *et al.* 2018). However, some estimates of ploidy were not consistent with prior information, therefore we used ratios of coverage obtained with Control-FREEC instead of inferred copy numbers. Control-FREEC was run using options *breakPointThreshold* = 0.8, *minExpectedGC* = 0.35, *maxExpectedGC* = 0.55, *telocentromeric* = 7000 and window size set to 250 bp. To estimate the ratio of coverage in maltose metabolic process genes (GO:0000023), we averaged coverage ratio for windows overlapping each gene. Strains with average genome coverage below 10x were excluded. The reference genome lacks some of the maltose genes, such as *MAL4* or *MAL6*, which are homologous to other *MAL* genes, and in case of their presence in the genome, could potentially affect read coverage of *MAL* genes. Although we cannot precisely estimate the number of copies of maltose metabolism genes, we can use our approach to roughly distinguish different categories of beer strains.

### Detecting introgression from *Saccharomyces* species

To detect potential gene flow between the Jean-Talon strain and other *Saccharomyces* species, we performed competitive mapping using SppIDER (last download 29/09/2019, (Langdon *et al.* 2018). In the first analysis, we concatenated genome assemblies of 8 *Saccharomyces* species: *S. paradoxus, S. cerevisiae, S. eubayanus, S. jurei, S. kudriavzevii, S. mikatae, S. uvarum* and *S. arboricolus.* In the second analysis, we combined genome assemblies of 6 lineages of S. paradoxus and the genome of *S. cerevisiae* S288C. All assemblies were masked using RepeatMasker v4.0.7 (http://www.repeatmasker.org) prior to the analysis.

### Assembly of the Jean-Talon genome

We assembled the Jean-Talon genome using the ONT dataset. We used wtdbg2 v2.5 (Ruan and Li 2019) with parameters -x ont -g 12m. The ONT reads were mapped against the draft assembly using minimap2 v2.17 (Li 2018) with parameter -x map-ont. The draft assembly was then polished using Nanopolish v0.11.1 (Loman *et al.* 2015) with parameter --min-candidate-frequency 0.1. Illumina reads were mapped against the signal-level polished assembly using bwa mem v0.7.16 (Li 2013) and the alignment was used to further polish the assembly using Pilon v1.22 (Walker *et al.* 2014). The polished assembly was aligned to the S288c reference genome from Yue et al. (Yue *et al.* 2017) using Mauve v2.4.0 (Darling *et al.* 2010), following which contigs were reordered to match reference chromosomes. Contig ctg6_pilon was manually split at the Ty2 junction as our structural analysis provided no support for the assembled translocation (see below). Visualization of translocations were produced with the Mauve GUI. The assembly is available at NCBI (PRJNA604588).

### Simulation of translocations in the S288c genome

The S288c genome was used to simulate three reciprocal translocations. Our goal was to estimate the power of a split mapping approach to detect rearrangements occurring at full-length Ty retrotransposon loci, since those are large (∼6kb) dispersed repeats which are expected to produce non-unique mappings. Using the genome annotations of Yue *et al.* (Yue *et al.* 2017) (coordinates are shown in parenthesis), two pairs of same-strand, full-length Ty1 elements were selected. Translocations were simulated between members of a pair using a custom python v3.7.1 script. The first translocation is between a subtelomeric Ty1 on chromosome VIII (562107-568134) and a Ty1 on chromosome XIII (378473-384398). The second translocation is between Ty1s on chromosomes IV (1214433-1220350) and XIV (512688-518577). A third translocation with genic breakpoints was simulated between YER068W on chromosome V (293281-295044) and YJR010W on chromosome X (462156-463691). The rearranged assembly harboring the three translocations was used to simulate PacBio reads using PBSIM v1.0.4 (Ono *et al.* 2013) with parameters --data-type CLR --depth 600 --length-mean 3000 --length-sd 2300 and the default error model.

### Structural variants (SVs) analysis using long reads

ONT reads for Jean-Talon (this study), PacBio reads for A.2565, A.T-58 (Fay *et al.* 2019) and S288c (Yue *et al.* 2017) and simulated PacBio reads for S288c were filtered with SeqKit (Shen *et al.* 2016) to keep read lengths between 8kb and 20kb inclusively. The filtered reads were mapped on the S288c reference genome (Yue *et al.* 2017) using minimap2 v2.17 with parameters -x map-ont or -x map-pb. SVIM v1.1.1 (Heller and Vingron 2019) was used to call five classes of SVs (deletions, insertions, tandem or interspersed duplications, inversions) based on the long-read alignments. Since the coverage depth of the Jean-Talon library was higher (59X) than that of A.2565 (9X) and A.T-58 (12X), the Jean-Talon library was randomly subsampled to approximately 9X using seqtk (https://github.com/lh3/seqtk) to correct for potential coverage depth biases in the detection of SVs. SVs supported by a number of reads lower than 15% of the coverage depth were filtered out. For each strain, we derived the distribution of physical distances from an SV call to the closest SV call of the same class in each of the other strains. Using the distribution of distances between the two Jean-Talon datasets (59X and 9X coverage depth) as a reference, we used one-sided Mann-Whitney U tests to determine which distributions were significantly shifted towards larger distances. Interspersed duplications and inversions were excluded from this analysis, as they comprised few or no calls across the datasets.

### Translocation analysis using split mapping

Long-read split mappings were used to search for translocations in the Jean-Talon, A.2565 and A.T-58 genomes compared to S288c. From the previously described long-read alignments, we extracted read IDs which had supplementary mappings and no secondary mapping using samtools v1.9 (Li *et al.* 2009). The alignments were filtered according to these read IDs using picard FilterSamReads v2.18.5 (http://broadinstitute.github.io/picard/) and subsequently analyzed using custom python v3.7.1 scripts. Keeping only the split reads which map to exactly 2 different chromosomes, we binned them in 20kb non-overlapping windows and represented read segment mappings with heatmaps. The length of the supporting reads was used to convert counts of supporting reads into approximate fraction of genome-wide coverage depth. Mappings of the S288c PacBio dataset against the reference S288c assembly allowed to identify artefactual signals arising from the split mapping approach. Mappings of the S288c simulated PacBio dataset against the reference S288c assembly allowed to compare the power of the split mapping approach to detect translocations at Ty and genic breakpoints.

## Results and Discussion

The colonies grown on YPD medium were creamy-beige round colonies, convex elevation, and matt finish, quite smooth and creamy surface. The diameter of most colonies is around 3 mm (Supplementary Figure 1). Under the microscope the cells looked round, medium and uniform in size and shape and arranged in clusters (Supplementary Figure 1), consistent with being *S. cerevisiae*.

### The Jean-Talon strain is a tetraploid strain and largely sterile

We first examine the ploidy and ability to sporulate of the Jean-Talon strain. DNA staining of 90 isolated colonies shows that it is a tetraploid (Figure 1A). Tetraploidy is also suggested based on the observed frequency distributions of single nucleotide polymorphisms (SNPs) mapped to the S288C genome that show peaks around frequencies of 0.25, 0.5 and 0.75 (Figure 1B). A recent study of the genome of 1011 *S. cerevisiae* isolates revealed that most of the natural isolates are diploid (Peter *et al.* 2018). However, approximately 11.5% of isolates are polyploid (3–5*n*) and those are enriched in specific subpopulations such as the beer, mixed-origin and African palm wine clades, which strongly suggests that some human-related environments have had an effect on the ploidy level (Peter et al., 2018). Similar results from (Gallone *et al.* 2016) and (Gonçalves *et al.* 2016) showed that multiple populations of beer strains show high rates of tetraploidy. Although spontaneous yeast tetraploids are usually fertile (Charron *et al.* 2019), the Jean-Talon strain shows about 30% spore viability (Figure 1C) and is close to the average spore viability observed for beers from the genetic groups Beer2, Mixed and Mosaic in (Gallone *et al.* 2016). The Jean-Talon is therefore typical of beer strains with respect to ploidy and fertility.

**Figure 1.**
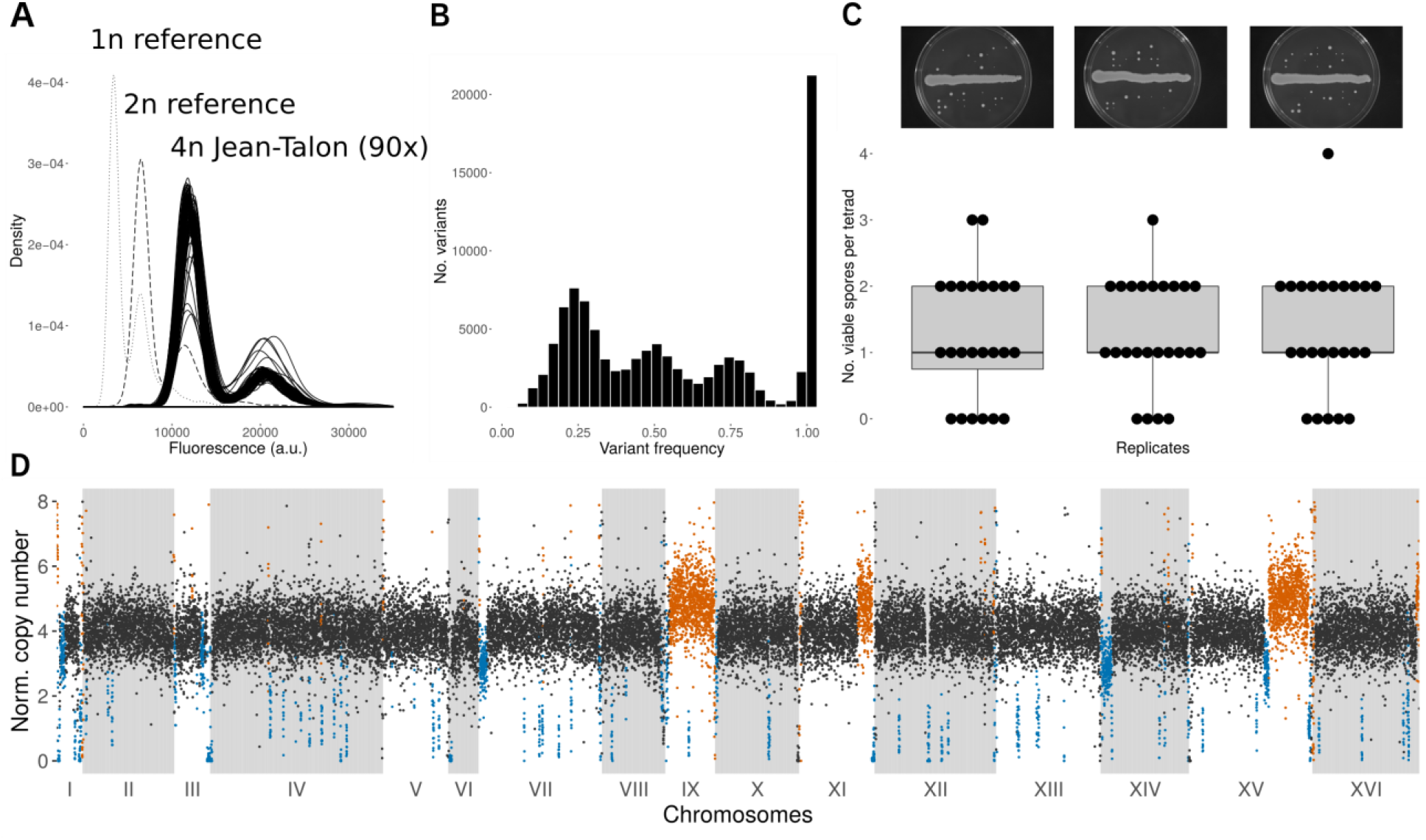
The Jean-Talon strain is a tetraploid with reduced spore viability. (A) DNA staining and fluorescence measured for 90 colonies of the Jean-Talon strain shows peaks around the expected ploidy of 4*n*. (B) Frequency of SNPs of the Jean-Talon strain reads mapped to *Saccharomyces cerevisiae* S288C genome. (C) Boxplots and numbers of viable spores per tetrad found in 3 biological replicates of sporulated cultures. Pictures of the spore dissection plates for the corresponding replicates are shown above. (D) Copy number profile up to 8*n* of the genome of Jean-Talon strain measured in 250 bp windows. Black windows correspond to the expected ploidy 4*n*, orange windows correspond to copy number gains and blue windows to copy number losses. Long stretches of copy number gains and losses correspond to ploidy of 5*n* and 3*n*, respectively.

Several long-range copy number gains and losses were observed in the genome, including the presence of 5 copies of chromosome IX, similar copy number changes at the ends of chromosomes XI and XV, and 3 copies at the beginning of chromosomes I, VII, XIV and middle of chromosome XV (Figure 1D). The aneuploidies and CNVs are typical of what is observed for industrial yeast strains (Gallone *et al.* 2016), but that are rare in species that have not been domesticated (Leducq *et al.* 2016) (Yue *et al.* 2017).

### The Jean-Talon strain belongs to the Beer/baking beer group

To find out to which genetic group Jean-Talon belongs to, we combined SNPs of the Jean-Talon strain with two yeast datasets: 401 strains from Fay et al. (Fay *et al.* 2019), and 1011 strains from 1000 yeast (Peter *et al.* 2018). Principal Component Analysis (PCA) on the Fay et al. dataset shows that Jean-Talon groups with the beer strains from the Beer/baking group according to PC2 and PC3 (Figure 2A), whereas PCA on the 1000 yeast dataset shows Jean-Talon grouping with the Mixed origin group, according to PC7 and PC8 (Figure 2B). The Mixed origin and Beer/baking groups comprise strains obtained from bakeries, breweries, as well as strains found in nature. Because the Jean-Talon strain was isolated from the environment, it may have mixed with other species, particularly *S. paradoxus*, which is found in Northern parts of North America (Charron *et al.* 2014) and with which it was shown to hybridize in different contexts (Barbosa *et al.* 2016). We did not detect gene-flow between the Jean-Talon and other *Saccharomyces* species using competitive mapping (Supplementary Figure 2).

**Figure 2.**
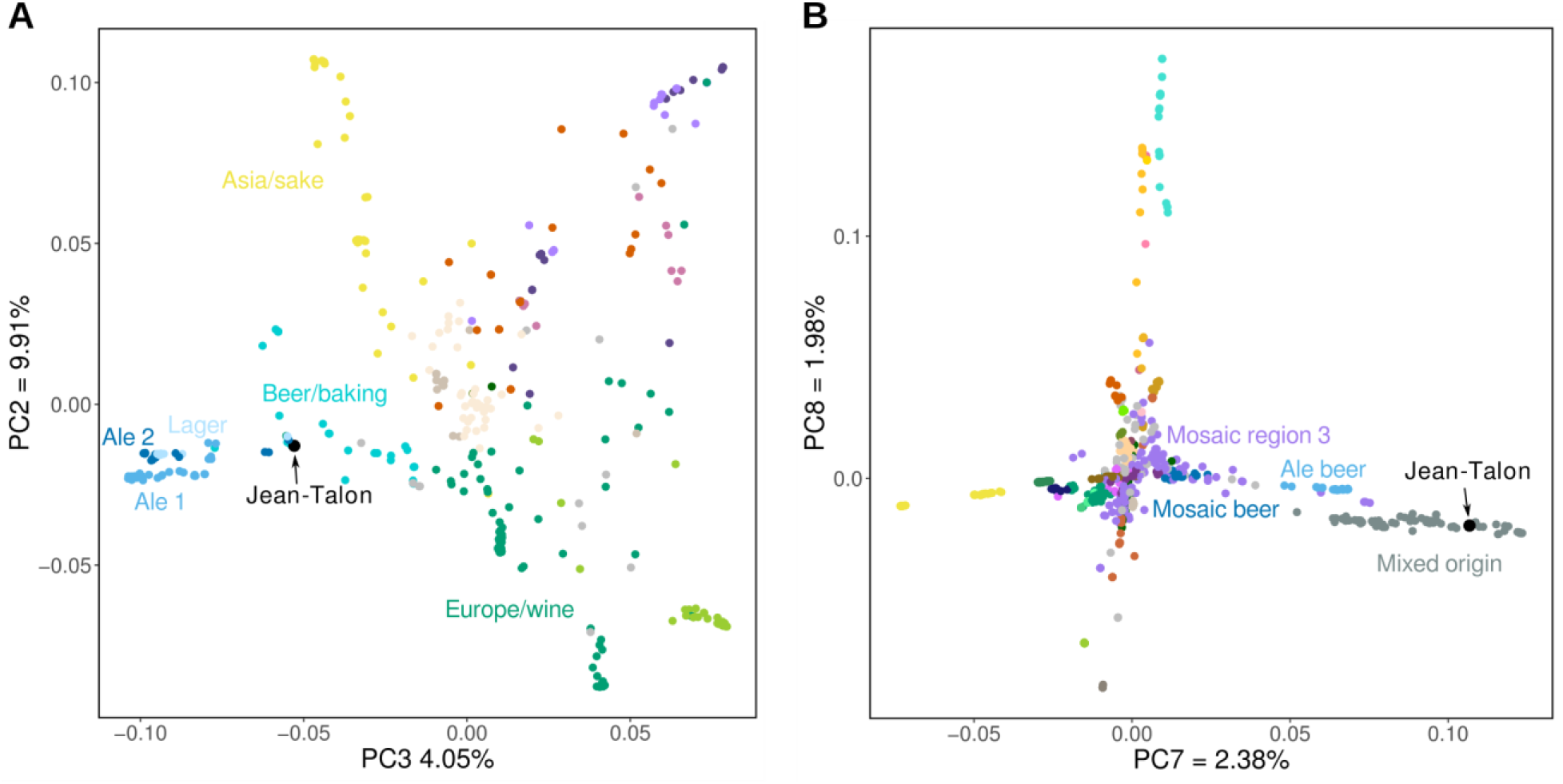
SNPs suggest that Jean-Talon belongs to the Beer/baking beer group. (A) PCA with 273,955 biallelic SNPs from 402 yeast strains from Fay et al. dataset groups Jean-Talon within the Beer/baking group. (B) PCA with 1,545,361 biallelic SNPs from 1000 yeast dataset (1013 yeast strains) groups Jean-Talon within the Mixed origin group.

To further investigate the Jean-Talon strain and identify the most closely related beer strains, we mapped the reads of 318 strains from 4 different studies, which include major beer groups (Supplementary Table 1) (Gallone *et al.* 2016; Gonçalves *et al.* 2016; Peter *et al.* 2018; Fay *et al.* 2019). Based on genotype similarity, the Jean-Talon strain is located on a branch composed of commercial beer strains (Figure 3A). According to the kinship coefficients, the strain is nearly identical (kinship coefficient between 93% and 97%) to the six other beer strains (Figure 3B), which include ales from England and Belgium (Supplementary Table 1). Estimates of synonymous heterozygosity and pairwise divergence between these strains support the finding that most segregating variants in the Jean-Talon are shared with other strains (Figure 3C). Using molecular clock, the time of divergence between Jean-Talon and S288C reference genome strain is about 14.4 M generations. Assuming constant mutation rate, the time of divergence of Jean-Talon from the closely related strains is equal to a fraction of 6.2E-05 of divergence time with S288C, which translates to 708 to 1080 generations, depending on the strain. The suggested number of generations per year in domesticated and lab yeast ranges from 150 (Gallone *et al.* 2016) to 2920 per year (Fay and Benavides 2005), however the growth of Jean-Talon could have been impeded when it stayed in the vaults. Moreover, generation time can be overestimated if breweries use the same yeast stock for each batch of fermentation, instead of continuously transferring yeast from one fermentation to the next. Although we do not know for how long the strain was dormant within the vaults, a small number of generations suggests that the split with other beer strains has not occurred long time ago. It is likely that the strain was used in the Boswell brewery which was still active in the 60s of the 20th century. Strains related to Jean-Talon were sampled from commercial ales, brewed from general purpose common yeast strains used for brewing different styles of beer, therefore they could have originated in the large commercial brewery.

**Figure 3.**
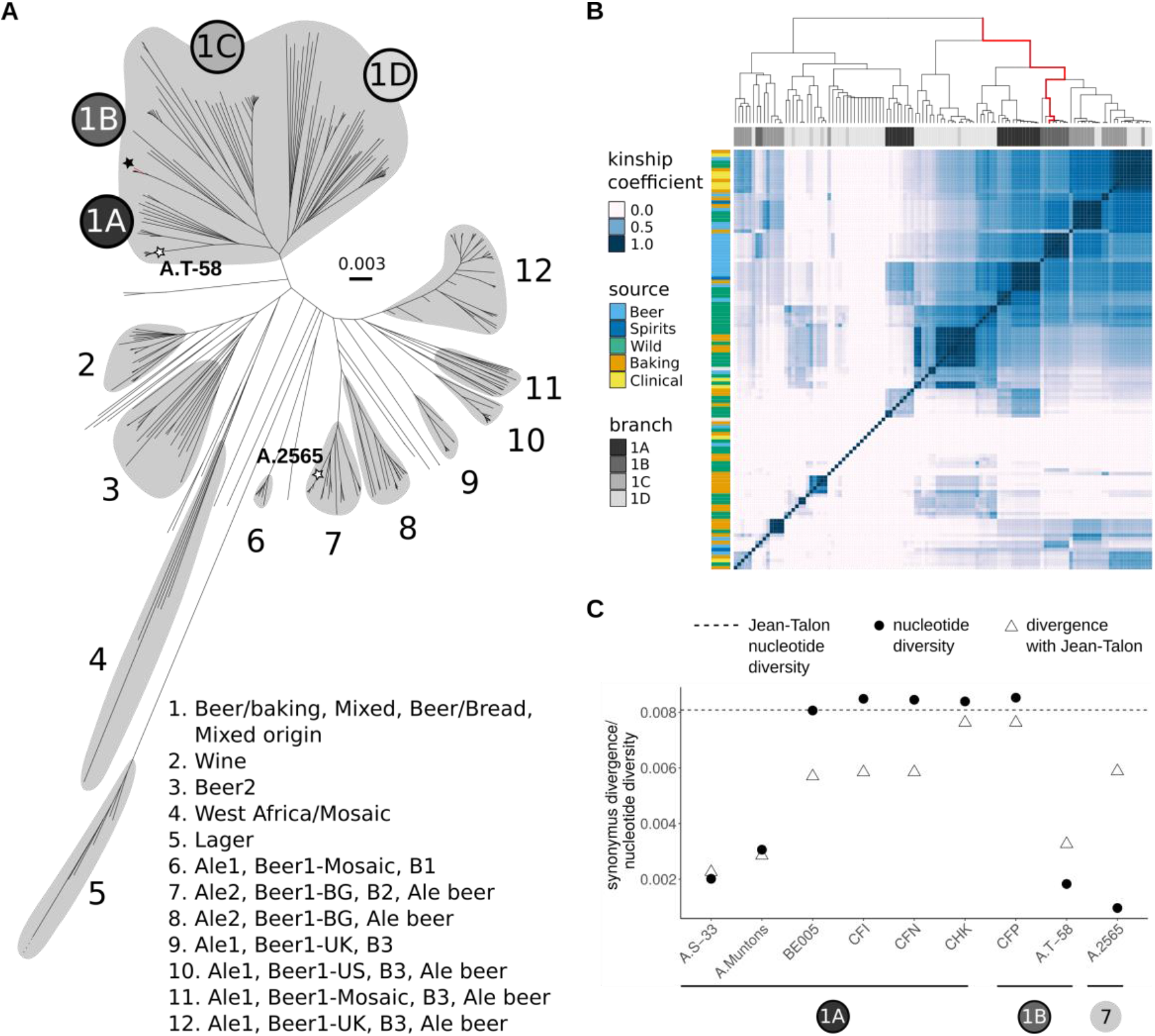
Jean-Talon is closely related to several commercial beer strains from the Beer/baking group. (A) Neighbor joining tree based on genome-wide genotype dissimilarity matrix for beer yeast strains from the four studies, Fay et al. 2018 (Ale1, Ale2, Lager, Beer/baking), Gallone et al. 2016 (Beer1*, Beer2, Mixed, Wine, West Africa, *Mosaic), Goncalves et al. 2016 (B1, B2, B3, Beer/Bread) and Peter et al. 2018 (Mixed origin, Ale beer). Consecutive numbers describe group affiliation of larger branches according to different studies. Note that one longest branch of the lager strain was cut to fit in the figure (dotted lines). The strain of Jean-Talon within the Beer/baking group is depicted with a black star, white stars depict the location of two beer strains with long-read sequencing data. (B) Heatmap of kinship coefficients estimated for all pairs of Beer/baking strains with 131,808 genome-wide SNPs. The Jean-Talon strain (red line on a dendrogram) has a kinship coefficient above 93% with 6 beer strains: CFI, CFN, BE005, A.Muntons, A.S-33, and A.Windson. (C) Nucleotide diversity in Jean-Talon strain is higher than divergence between most closely related beer strains, suggesting that many segregating variants are shared between the strains. Circles with numbers depict tree branches from (A).

### Distinct structural variation of the Jean-Talon strain

The profiles of copy number variation across the genomes of the related strains show multiple aneuploidies, mostly shared with the Jean-Talon strain, supporting their recent divergence (Supplementary Figure 3). The important exception is a 350 kb region with 5 copies located on chromosome XV. We examined specifically the copy number of genes that have been associated with adaptation of beer strains to the brewing environment, for instance maltose metabolism genes. Hierarchical clustering of the beer strains based on the number of copies in genes involved in maltose metabolic process groups Jean-Talon and its related strains with undefined ale beers from Belgium and England and lagers (Figure 4). We called five classes of structural variants (SVs) based on mappings of long reads to the S288c reference genome (Figure 5A). Distributions of physical proximity of SV calls between strains show that Jean-Talon is closer to a strain from the Beer/baking group (A.T-58), rather than to a strain from the Ale2 group (A.2565, Figure 5B). Despite this relatedness, the Jean-Talon strain exhibits a distinct pattern of SVs as it is significantly different from both beer strains for the most abundant classes of SVs (insertions and deletions, Figure 5B).

**Figure 4.**
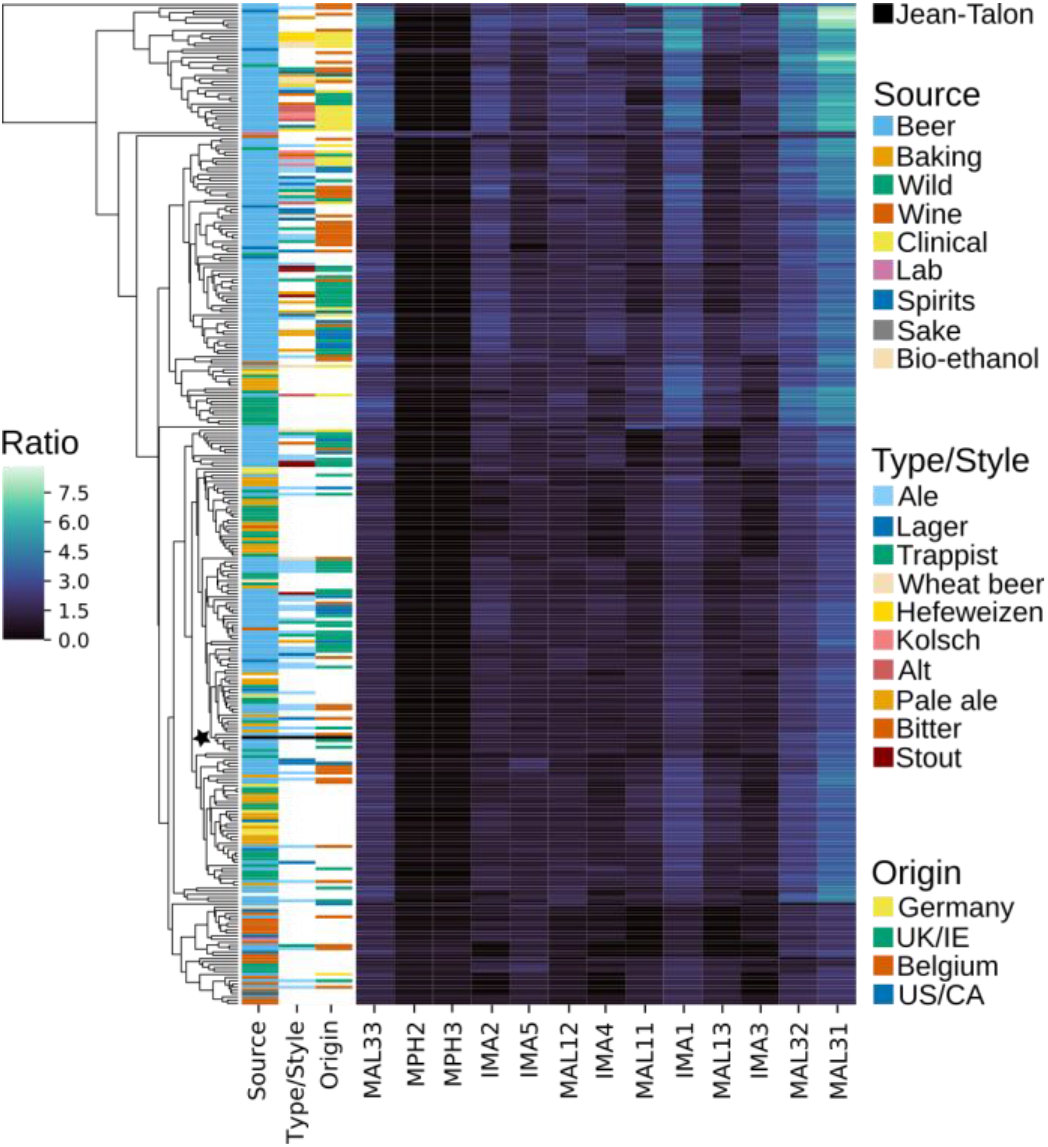
Clustering of copy number variation in genes involved in maltose metabolic processes place Jean-Talon among Belgium and English ales and lagers. Ratio indicates copy number change relative to the strain ploidy. Beer styles, types and origin differentiate in the matrix mainly according to changes in copy number in *MAL3* and *IMA1* genes. Black star and line depict location of the Jean-Talon strain.

**Figure 5.**
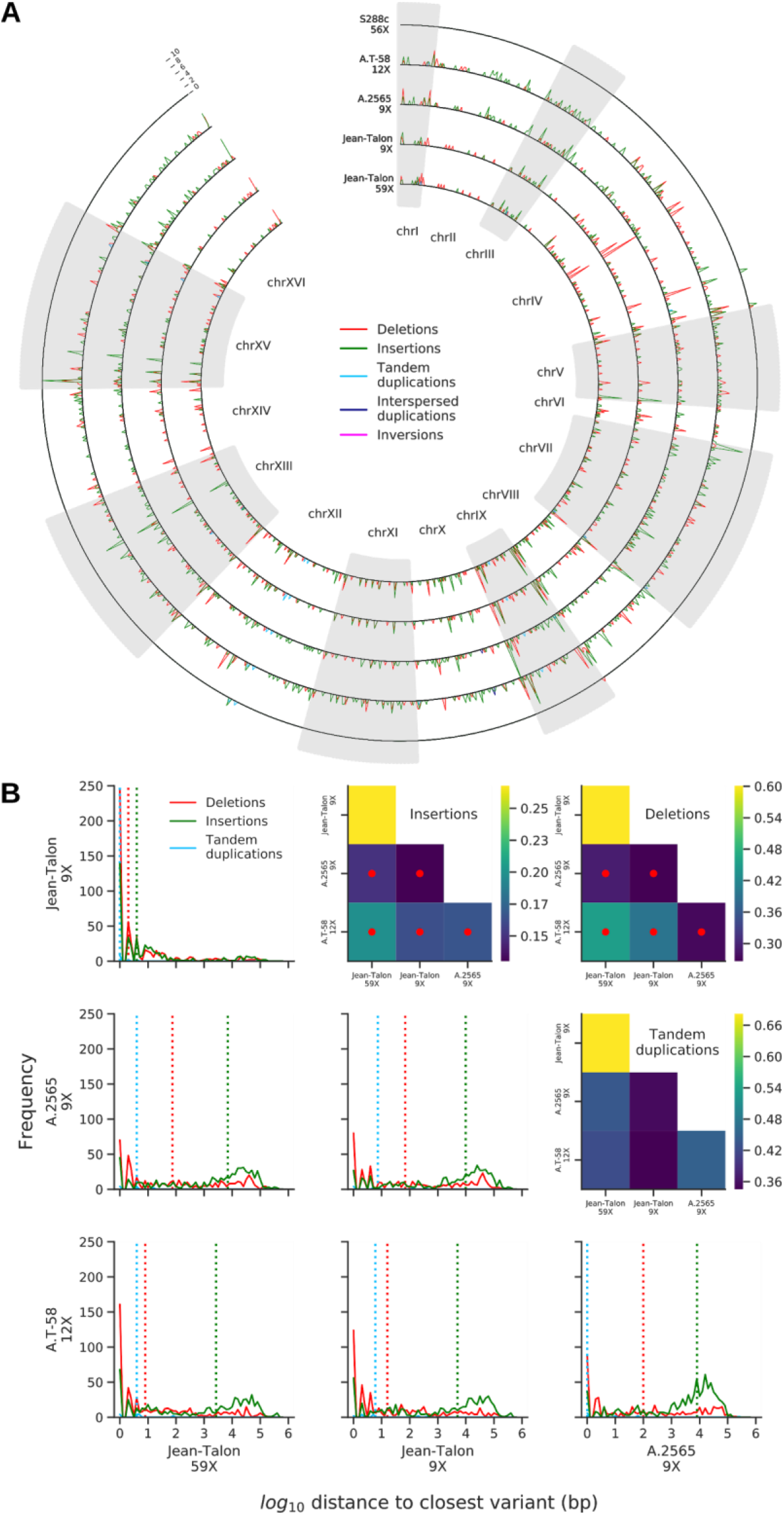
Structural variation in the genome of the Jean-Talon strain. (A) SVs against S288c for the Jean-Talon strain (complete and subsampled datasets), and two beer strains with available long read datasets, one from the Beer/baking group (A.T-58) and one from the Ale2 group (A.2565). SV density in non-overlapping 10 kb windows is plotted. (B) Physical proximity of SV calls between strains. Distributions of physical distance to the closest same-class SV call in the mate strain is shown for each pair of strains. Dotted vertical lines correspond to medians. Heatmaps show the results of two-sided Mann-Whitney U tests for each distribution compared to the (Jean-Talon 9X: Jean-Talon 59X) reference pair. Color maps show ratios of U statistics to the reference, while red dots indicate distributions significantly right-shifted compared to the reference (p-values<0.05, Mann-Whitney U tests, FDR corrected).

We assembled the Jean-Talon genome using our Oxford Nanopore dataset and detected a translocation between chromosomes II and XI (Supplementary Figure 4), which breakpoint maps to a full-length Ty2 retrotransposon. We used split mappings of long reads to investigate translocations in the Jean-Talon, A.T-58 and A.2565 strains compared to S288c (Supplementary Figure 5). Although we found this method has reduced power to detect translocations at Ty loci compared to genic breakpoints, we find no evidence for a translocation between chromosomes II and XI in Jean-Talon (Supplementary Figure 6). Thus, this translocation is likely an assembly artefact and the genomes of Jean-Talon and S288c appear to be collinear.

## Conclusion

The yeast Jean-Talon strain was isolated from an archeological site in the old part of Québec City where the first brewery was founded in the 17th century. The strain was isolated from the vaults of the second Intendant’s palace that was built in the 18th century and occupied by the Boswell brewery starting in the 19th century. The Jean-Talon strain is a strain of *Saccharomyces cerevisiae*, which is not found naturally in this part of North America (Charron *et al.* 2014). The strain is very closely related to other strains used in industrial brewing, suggesting that it derived recently from other industrial beer strains.

## Acknowledgements

We thank members of the Landry lab for input on the project. This work was funded by a Natural Sciences and Engineering Research Council of Canada (NSERC) Discovery grant to CRL. AF was supported by a Genome Canada Grant (LSARP BIOSAFE), MH was supported by a NSERC Alexander Graham Bell graduate fellowship and SM by a Fonds de recherche santé Québec (FRSQ) postdoctoral fellowship. CRL holds the Canada Research Chair in Evolutionary Cell and Systems Biology.

## Data availability

Raw short and long sequencing reads and the genome assembly are available at NCBI (PRJNA604588).

