## Supplementary Figures for "The genome sequence of the Jean-Talon strain, an archeological tetraploid beer yeast from Québec"

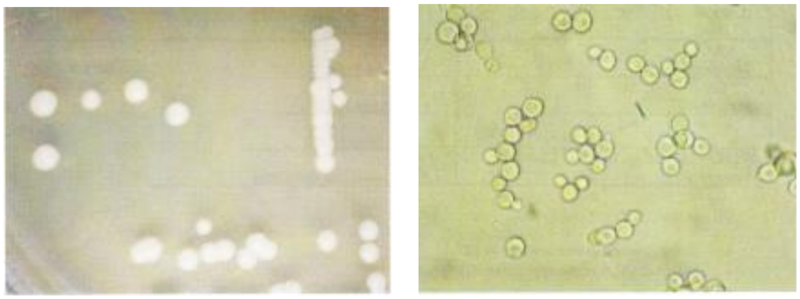

**Supplementary Figure 1. Colonies and cells of the Jean-Talon strain.** On the left picture, colonies on YPD media, after 48h incubation at 30°C. On the right picture yeast cells in YPD, after 16h incubation at 28°, 175 rpm, observed at 40X.

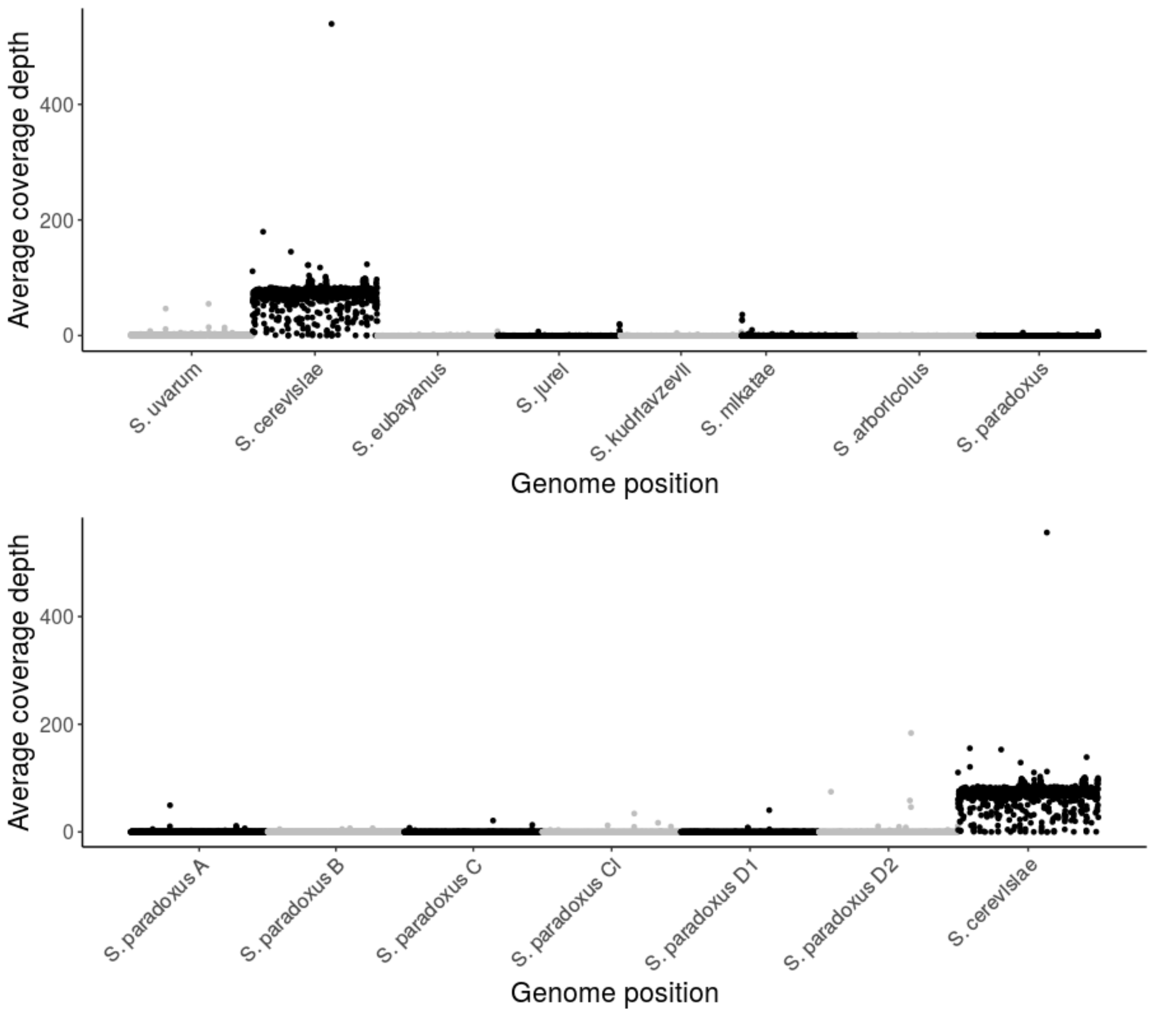


**Supplementary Figure 2. No evidence for gene flow from/to *Saccharomyces* species or *S. paradoxus*.** Coverage depth of Jean-Talon reads mapped to concatenated genomes of 8 *Saccharomyces* species (top) and concatenated genomes of *S. cerevisiae* and 6 *S. paradoxus* lineages (bottom). Reads map exclusively to the *S. cerevisiae* genome. Coverage depth in the plot was averaged across 10 kb non-overlapping windows. Statistically significant positive residuals of a $\chi^2$ test of randomness of read counts across all genomes (p-value <2.2E-16) are produced only by *S. cerevisiae*.

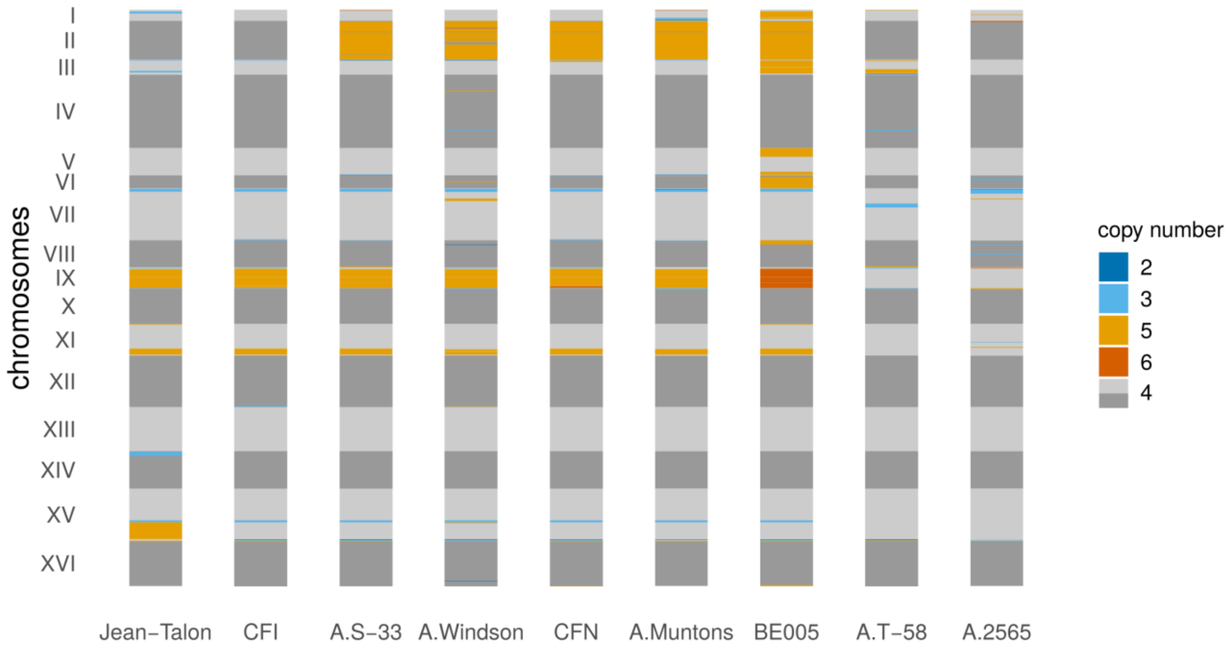


**Supplementary Figure 3. Jean-Talon shares most aneuploidies with other strains in the Beer/baking group.** We show copy number profiles for regions of at least 10 kb, in which > 90% of 250 bp windows have a given copy number. Jean-Talon has 5 copies of a 350 kb long fragment on chromosome XV which is present in 4 copies in other beer strains.

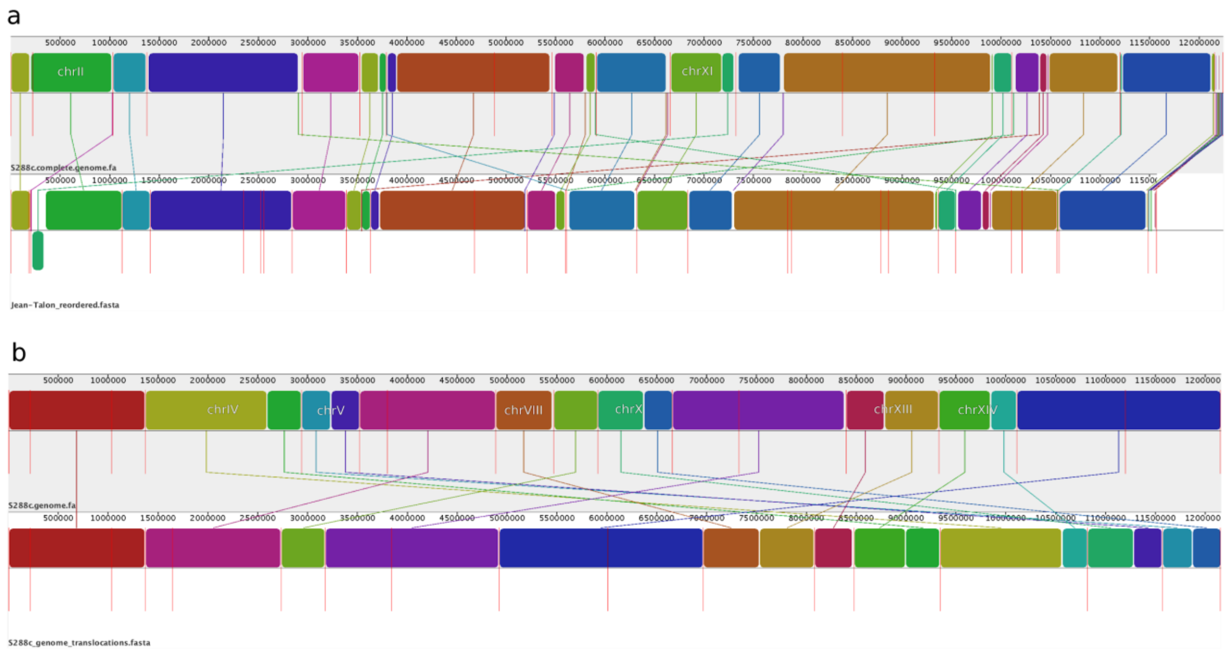


**Supplementary Figure 4. Illustration of translocations from genome assembly and simulations.** (A) Translocation between the right arm of chromosome XI and the left arm chromosome II in the Jean-Talon long-read assembly. The assembled translocation junction corresponds to a full-length Ty2 retrotransposon. Upper track is the S288c reference assembly from Yue *et al.* 2017; lower track is our Jean-Talon assembly. (B) Translocations simulated from the S288c reference assembly (upper track). Lower track: translocations between chromosomes IV and XIV and between chromosomes VIII and XIII have breakpoints at full-length Ty1 retrotransposons, the Ty1 on chromosome VIII being subtelomeric; the breakpoint of the translocation between chromosomes V and X is within the genic sequences.

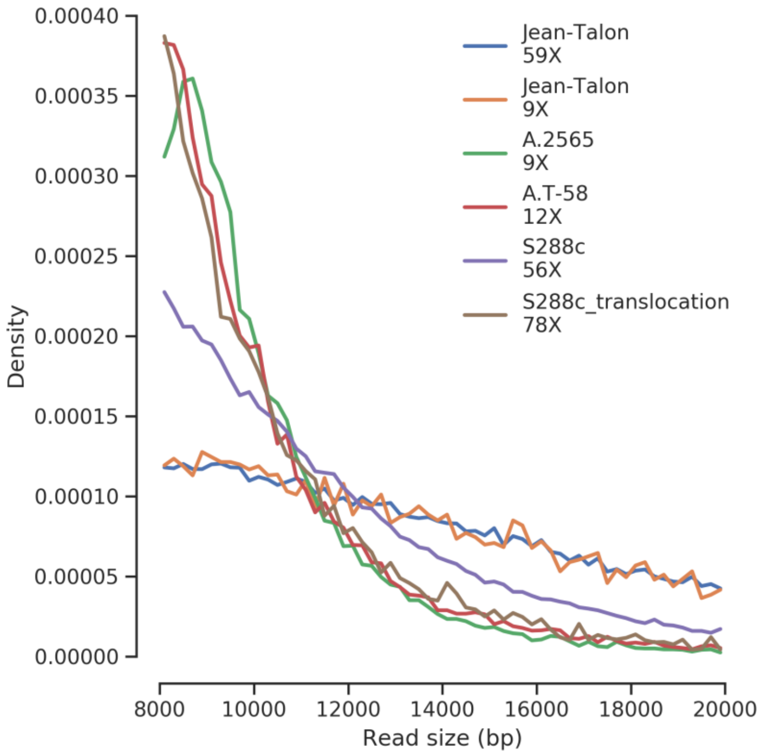


**Supplementary Figure 5. Read length distribution for the long-read datasets used in this study.** Jean-Talon corresponds to the Oxford Nanopore dataset generated in this study; A.2565 and A.T-58 correspond to PacBio datasets from Fay *et al.* 2019; S288c corresponds to the PacBio dataset from Yue *et al.* 2017; S288c_translocation corresponds to the simulated PacBio dataset from the S288c assembly harboring translocations as shown in Supplementary Figure 4.

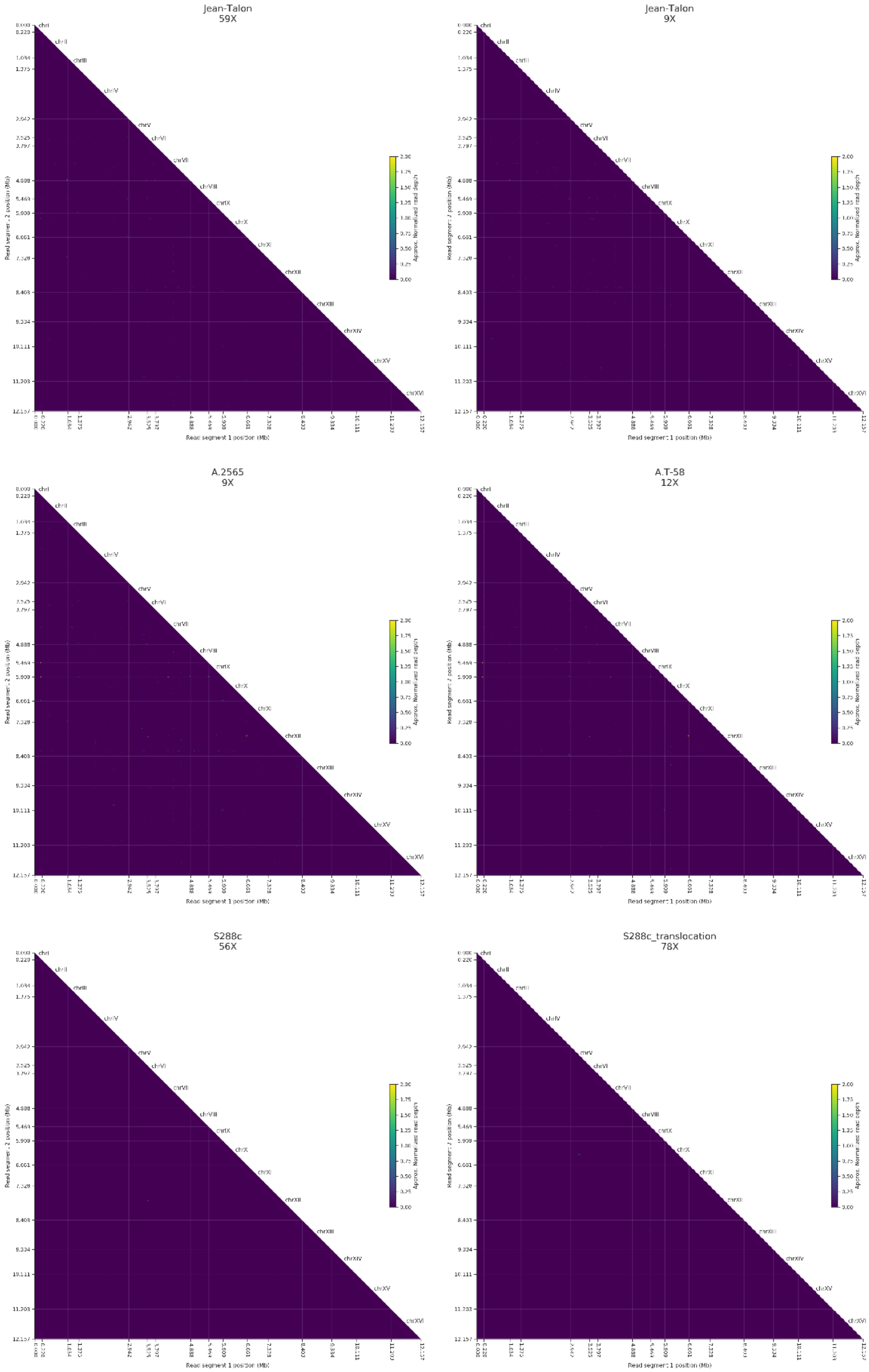
Jean-Talon
59X
Jean-Talon
9X
A.2565
9X
A.T-58
12X
S288c
56X
S288c_translocation
78X
chrI
chrII
chrIII
chrIV
chrV
chrVI
chrVII
chrVIII
chrIX
chrX
chrXI
chrXII
chrXIII
chrXIV
chrXV
chrXVI
Read segment 1 position (Mb)
Read segment 2 position (Mb)
Approx. Normalized read depth

**Supplementary Figure 6. Split mapping of long reads fails to detect the translocation between chromosomes XI and II in the Jean-Talon strain.** Heatmaps show the number of reads which have supplementary mappings to exactly two chromosomes in windows of 20kb. Color maps correspond to the approximate genome-wide coverage depth equivalent of the supporting reads (the upper bound corresponds to coverage depth equal or higher than 2). Simulated translocations (bottom right) show that breakpoints at Ty1 elements are difficult to detect with the split mapping approach (coverage depth equivalent <1), while the genic breakpoint is correctly detected (coverage depth equivalent around 1). Mapping of the S288c dataset against the S288c assembly (bottom left) illustrates some artefacts of the split mapping approach, where signal is detected in many other strains. Although we cannot exclude that some of the signal from the Jean-Talon strain (top) and other beer strains (middle) might not be artefactual, there is no signal corresponding to the translocation detected in the Jean-Talon assembly.
